# Structure of a fungal 1,3-β-glucan synthase

**DOI:** 10.1101/2023.03.19.532754

**Authors:** Chao-Ran Zhao, Zi-Long You, Dan-Dan Chen, Jing Hang, Zhao-Bin Wang, Le-Xuan Wang, Peng Zhao, Jie Qiao, Cai-Hong Yun, Lin Bai

## Abstract

1,3-β-Glucan is the major component of the fungal cell wall and is synthesized by 1,3-β-glucan synthase located in the plasma membrane, which is a molecular target of anti-fungal drugs echinocandins and the triterpenoid ibrexafungerp. In this study, we report the 3.0-Å resolution cryo-EM structure of *Saccharomyces cerevisiae* 1,3-β-glucan synthase, Fks1. The structure reveals a central catalytic region adopting a cellulose synthase fold with a cytosolic conserved GT-A type glycosyltransferase domain and a closed transmembrane glucan-transporting channel. We found that two extracellular disulfide bonds are crucial for Fks1 enzymatic activity. Structural comparison between Fks1 and cellulose synthases and structure-guided mutagenesis studies provided novel insights into the molecular mechanisms of the fungal 1,3-β-glucan synthase.

## Introduction

Fungal diseases are a growing threat to human health. Approximately 1.7 billion people have skin, nail, and hair fungal infections, and over 1.6 million mortalities occur each year because of serious fungal diseases(*1, 2*). Furthermore, fungal pathogens cause major losses to agricultural activities and food production and account for approximately 65% of pathogen-driven host extinctions(*3, 4*). There are currently five classes of clinically approved antifungal drugs: polyenes (amphotericin B), flucytosines, azoles (ketoconazole, itraconazole, fluconazole, voriconazole, and isavuconazole), echinocandins (caspofungin, micafungin, and anidulafungin), and the triterpenoids (ibrexafungerp)(*5-7*). Among them, the echinocandins and triterpenoids are the only two new classes of antifungal drugs introduced into the market over the last 40 years(*1, 5, 6*). These two classes inhibit 1,3-β-glucan synthase in the plasma membrane to block fungal cell wall synthesis(*5, 6*).

Fungal cell walls are essential for the viability, morphogenesis, and pathogenesis of fungi(*8*). Fungal cell walls are an attractive target for anti-fungal drug development because such a cellular structure is absent in human cells(*9*). Unlike the plant cell wall that’s primarily made of cellulose (1,4-β-glucan), the fungal cell wall is composed mainly of 1,3-β-glucan, 1,6-β-glucan, chitin, and mannoproteins(*10*). Most approved and developing anti-fungal drugs that target the cell wall function by inhibiting 1,3-β-glucan synthase(*9*). This enzyme catalyzes the formation of β(1→3) glycosidic linkages in 1,3-β-glucan by using Uridine Di-Phosphate-activated glucose (UDP-Glc) as the sugar donor and transport the glucan through the membrane (**Fig. 1a**). Fungal 1,3-β-glucan synthase is primarily encoded by *FKS1*, or *FKS2*(*11*). *FKS1* is an essential gene in most fungi, including Candida and Aspergillus species(*12*). Simultaneous knockout of *Saccharomyces cerevisiae FKS1* and *FKS2* is lethal(*13*). Many echinocandins resistant mutations have been identified on three hot spots of *FKS1*, indicating that echinocandins target Fks1(*14, 15*). In *S. cerevisiae*, Fks1 is expressed predominantly under normal conditions, whereas Fks2 expression is important during sporulation or growth under particular stressful conditions(*13, 16*). *S. cerevisiae* Fks1 shares ∼88% identity to Fks2, and 56%–86% identity to FKSs in major pathogenic fungi species, suggesting high similar overall structures and mechanisms for different fungal 1,3-β-glucan synthases (**Supplementary Fig. 1**). The 1,3-β-glucan synthase activity of Fks1 is regulated by Rho1, which is a Ras-like small GTPase(*17-21*). Deficiency in 1,3-β-glucan synthase activity was caused by *RHO1* mutants and restored by purified Rho1 in vitro(*17, 18, 20*).

**Fig. 1.**
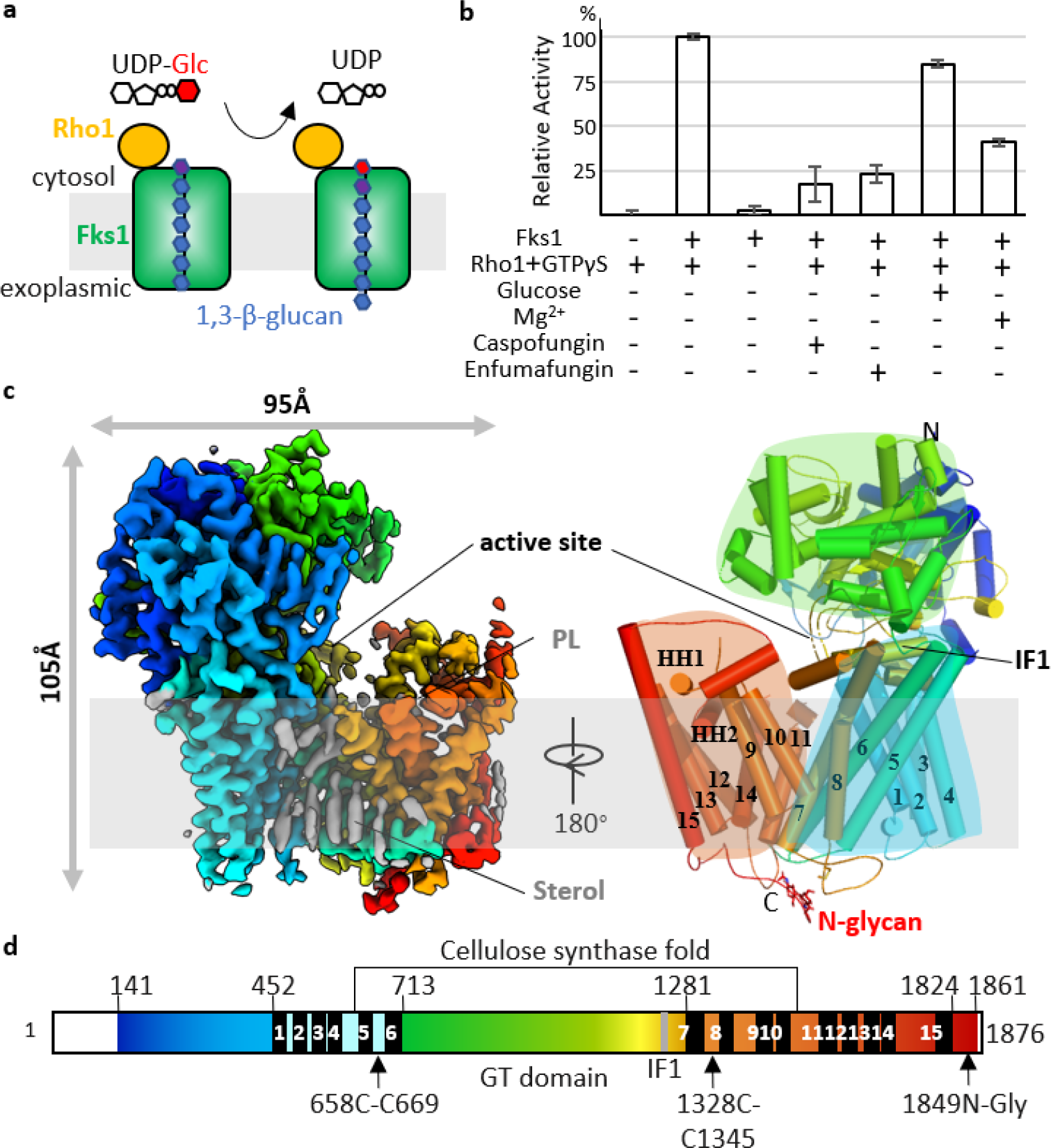
Enzyme activity and cryo-EM structure of Fks1. **a** Synthesis and transport scheme of 1,3-β-glucan by Fks1. The activity was found to be regulated by Rho1. **b** In vitro UDP-Glo glycosyltransferase assay with purified Fks1 with or without purified Rho1 and GTPγS. The assay under condition with Rho1 and GTPγS was also performed with glucose, Mg^2+^, caspofungin, or enfumafungin. The reaction without Fks1 was used as the control. Data points represent the mean ± SD in triplicate. **c** Cryo-EM map and atomic model of Fks1. The polypeptide is shown in rainbow from N- to C-terminus. **d** The Fks1 domain map. Major domains and motifs are labeled. The regions not observed are in white.

Fks1 is a GT-A type glycosyltransferase that belongs to the glycosyltransferase 48 (GT48) family (http://www.cazy.org). Despite its importance in cell wall synthesis and anti-fungal drug development, the structure of Fks1 has not been determined, hindering a mechanistic understanding of 1,3-β-glucan synthesis. In this report, the cryo-EM structure of *S. cerevisiae* Fks1 was solved at 3.0 Å resolution, which is the first structure of a GT48 family member. The structure provides insights into the catalytic mechanism and transport of glucan by Fks1 and how echinocandins inhibit the activity of this enzyme.

## Results

### Purification and characterization of Fks1

Fks1 was overexpressed in *S. cerevisiae* using a multicopy plasmid with a strong GAL promoter and a C-terminal triple FLAG tag and purified with an anti-FLAG affinity column followed by size-exclusion chromatography (**Supplementary Fig. 2a, b**). Although previous studies have identified Rho1p as a co-purified regulatory subunit of the functional glucan synthase(*17-20*), our purified sample only contained Fks1 (**Supplementary Fig. 2b**). In agreement with previous findings, our subsequent in vitro UDP-Glo glycosyltransferase assay of the purified sample revealed that Fks1 exhibited minimal activity by itself, but was highly activated by the addition of purified Rho1 and GTPγS (**Fig. 1b, Supplementary Fig. 2d, e**). We also found the activity of Fks1 was inhibited by the echinocandin drug caspofungin and the triterpenoid enfumafungin, which is the precursor of ibrexgerp (**Fig. 1b**). Previous studies have revealed that these inhibitors specifically target on 1,3-β-glucan synthase (*5, 22*), suggesting the activity we detected was indeed caused by 1,3-β-glucan synthase.

Fks1 belongs to the GT-A superfamily of glycosyltransferases. Most GT-A fold glycosyltransferases are metal-dependent with a conserved metal coordinate DxD motif(*23*). However, the corresponding sequence in Fks1 is a DAN motif (residues 1102–1104), which has been shown to be essential for Fks1 function by a growth complementary assay (**Supplementary Fig. 1**)(*15*). To verify whether Fks1 activity is affected by metal ion, we measured the in vitro UDP-Glc hydrolysis activity of Fks1 in the buffer with or without Mg^2+^ (**Fig. 1b**). We found that the presence of Mg^2+^ is not required for or even decreased the activity of Fks1, which is consistent with previous studies (*24, 25*). Besides, the activity of Fks1 was not affected by glucose in the reaction buffer (**Fig. 1b**), which is also consistent with previous studies that the activity of 1,3-β-glucan synthase does not need glucose as a starting primer (*24, 25*).

### The overall structure of Fks1

We performed single-particle cryo-EM on purified Fks1 and obtained a cryo-EM 3D map at 3.0-Å resolution (**Supplementary Figs. 2f, 2g, 3**). The 3D map had excellent main-chain connectivity and side-chain densities except for some disordered loops (**Fig. 1, Supplementary Fig. 4**). We built an atomic model into the map and refined the model to good statistics (**Supplementary Table 1**).

Yeast Fks1 has 1876 residues, and the structure measures ∼105 Å tall and 95 Å wide (**Figs. 1c, d, Supplementary Fig. 1**). Fks1 can be divided into a transmembrane region and a large cytosolic region. The transmembrane region contains 15 transmembrane helices (TM) that can be grouped into two parts, TM1–8 and TM9–15, based on the packing arrangement of these helices. The TM9–15 and two amphipathic horizontal helices (HH1–2), as a whole, tilt away from the first eight TMs to generate a sizable cavity in the middle of the structure facing toward the cytosol. This structural feature likely facilitates the transport of the glucan product through the membrane.

The cytosolic region of Fks1 is composed of two parts: an N-terminal peptide (Met-141 to Phe-452) and the GT domain inserted between TM6 and TM7 (Ile-713 to Gln-1281) (**Figs. 1c, d**). The first 140 residues of Fks1 are largely disordered and invisible in our cryo-EM map. The whole cytosolic region folds into a 9-stranded *β*-sheet, which is sandwiched by two groups of *α*-helices. The cytosolic region is tilted relative to the membrane plane and packs onto the first eight TMs of Fks1, and this interaction is mainly mediated by an amphipathic interface helix (IF1) (Lys-1172 to Gly-1188) in the center. Such architecture forms a large horizontal groove corresponding to the enzyme active site, as described below.

In addition, we identified several ordered sterol molecules and an ordered phospholipid in the transmembrane region (**Figs. 1c, d, Supplementary Figs. 4, 5**). The sterol molecules are found to be around TMs and likely belong to the detergent CHS that was added during purification. Purified Fks1 in buffer without CHS forms aggregates predominantly, suggesting that the sterols help to stabilize the overall Fks1 structure (**Supplementary Fig. 2a–c**). The phospholipid molecule is embedded in a cavity surrounded by TM9, TM14, and HH2. Noticeably, its phosphate group directly hydrogen-bonds to the side chains of R1455, R1684 and H1688 (**Supplementary Fig. 5**). This phospholipid probably stabilizes the C-terminal region of Fks1 and plays a structural role.

A structure-based homology search using the online Dali server indicated that the central part of Fks1 (Gly-615 to Asn-1490) shares a structural fold with bacterial cellulose synthase BcsA and plant cellulose synthase CesA in the GT-2 family (**Figs. 1d, 2a, 3a, Supplementary Fig. 6**)(*26-32*). Specifically, this part of Fks1 includes a cytosolic GT domain and a glucan transmembrane transport domain. Cellulose is a linear D-glucose chain with β(1→4) glycosidic linkages. Both Fks1 and cellulose synthases catalyze glycosidic bond formation using UDP-Glc as the sugar donor and transport the glucan through the membrane. Therefore, we performed structural alignments between Fks1 and BscA or CesA in complex with substrates (**Figs. 2–3, Supplementary Fig. 6**)(*26, 28*), and identified the putative catalytic site and glucan-transporting channel of Fks1. Because no substrate density was observed in catalytic site or glucan-transporting channel of Fks1, our Fks1 structure no doubt represents an apo form.

**Fig. 2.**
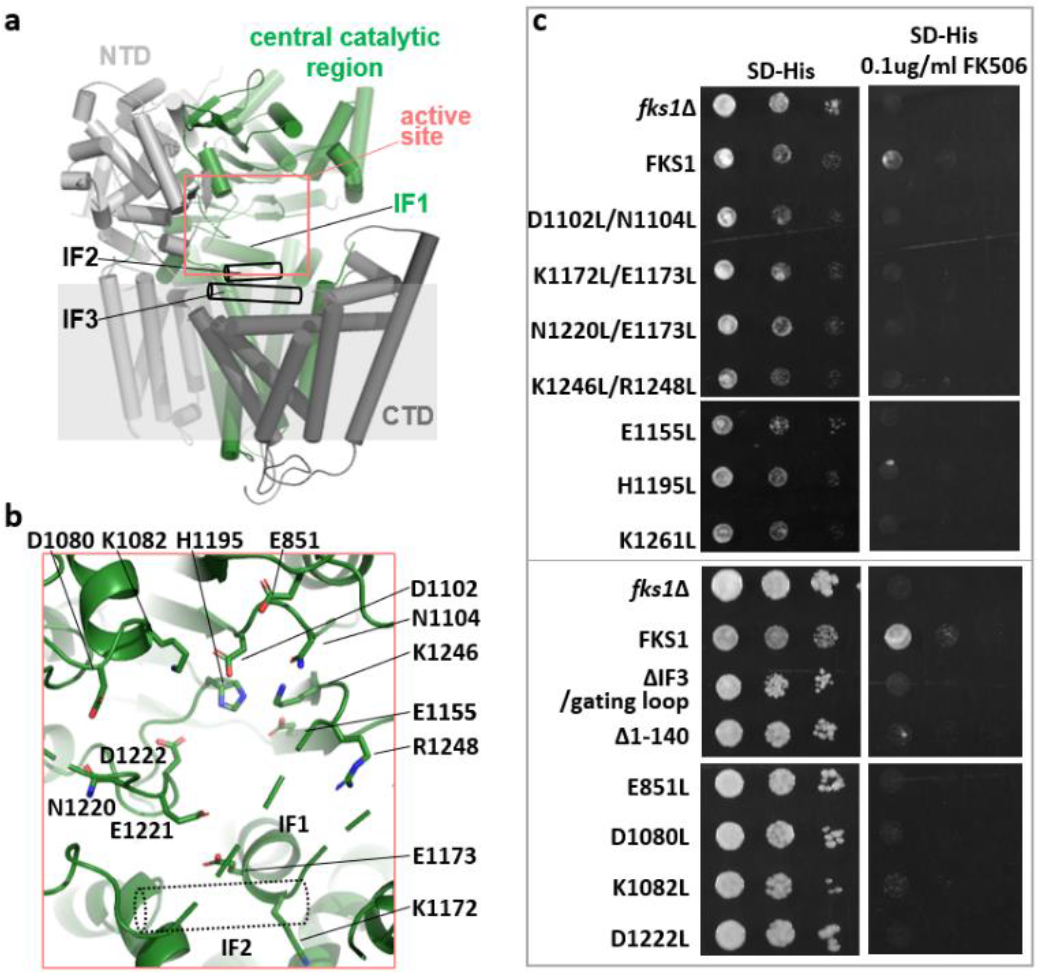
Putative active site in the GT domain of Fks1. **a** Overall structure of Fks1 in cartoon representation. The central catalytic region (green) is sandwiched by N-terminal (grey) and C-terminal (black) regulatory domains. The putative active site is highlighted by a circle (salmon). Flexible IF2 and IF3 are highlighted by two open black cylinders. **b** A putative active site in Fks1, with residues forming the pocket shown in stick representation. **c** Growth complementation of *fks1*Δ cells with empty plasmid (*fks1*Δ), or plasmid carrying either wild-type Fks1 (FKS1) or mutants. 2*μ*l (top box) or 5*μ*l (bottom box) cells were spotted onto synthetic Histidine-dropout medium plates (SD-His) and SD-His plates with 0.1 *μ*g/mL FK506. Plates were incubated at 30 °C for two days.

**Fig. 3.**
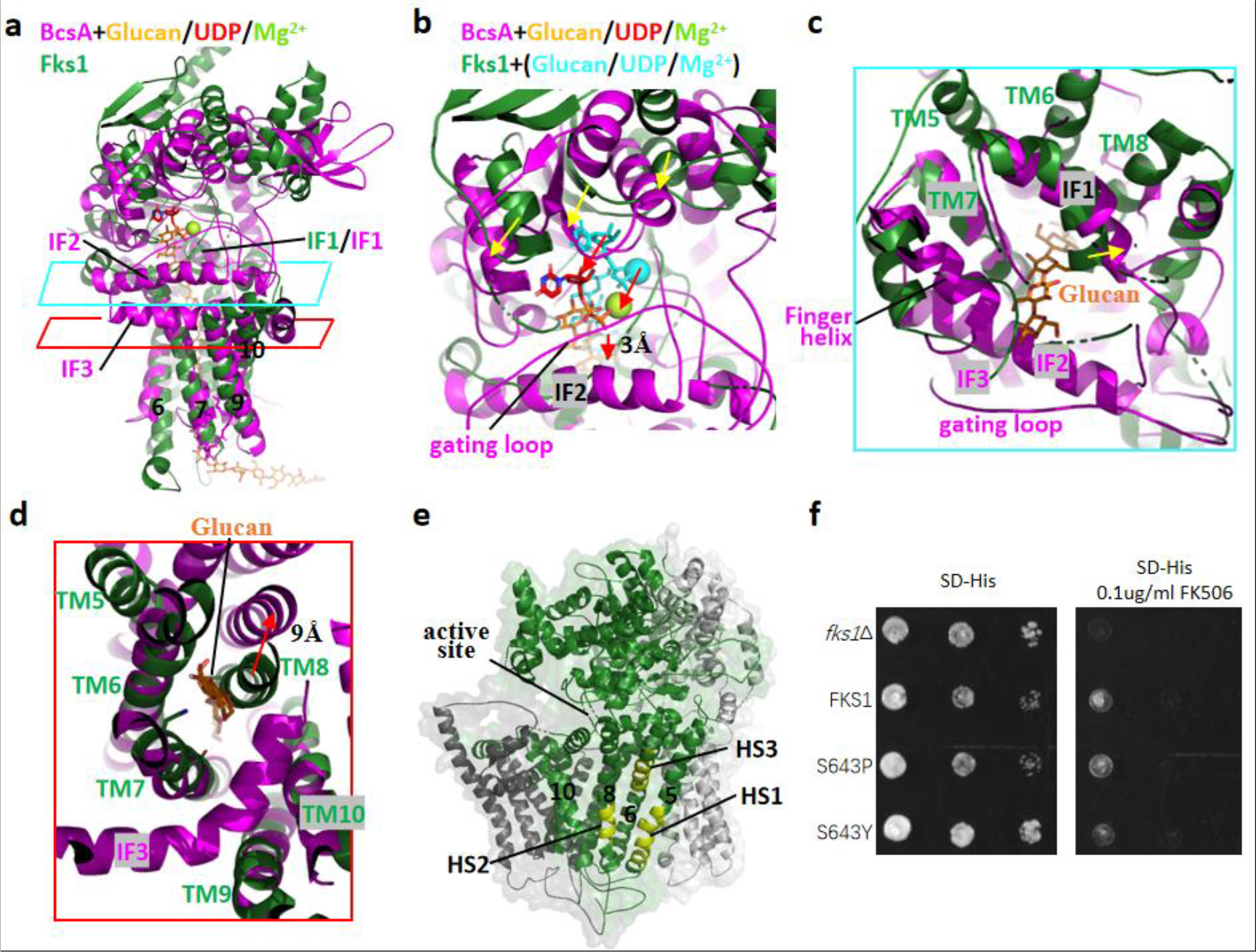
Putative glucan transporting channel of Fks1. **a** Structural comparison between the central catalytic region of Fks1 (green) and BcsA (PDB ID: 5EJZ, magenta) in complex with cellulose (orange), UDP (red) and Mg2+ (lemon) by aligning their respective TMs. The cyan and red rhomboids mark the positions for cut-in views shown in panels **c** and **d. b** A close-up view of the cytosolic region in panel **a**. Substrates in cyan are modeled onto Fks1’s active site basing on structural alignment of GT domains of Fks1 and BcsA in **Fig. 2b**. Movements of the GT domain and substrates are highlighted by yellow and red arrows, respectively. The glucan shifts downward by ∼3Å. **c, d** Cut-in views from the cytosolic side of the superposition of Fks1 and BcsA in panel **a**. TM8 and IF1 of Fks1 form part of the putative glucan transport path. Movements of the TM8 and IF1 are highlighted by red and yellow arrows, respectively. Compared with BcsA, TM8 of Fks1 is shifted inward by ∼9Å and occupy the glucan transporting path, resulting in a closed channel. Corresponding sequences of IF2–3, and gating loop are disordered in Fks1. **e** Mapping of the echinocandins resistance mutant hot-spots (HS1–3) onto the structure of Fks1. **f** Growth complementation of *fks1*Δ cells with empty plasmid (*fks1*Δ), or plasmid carrying either wild-type Fks1 (FKS1) or mutants (S643P and S643Y in HS2). Cells were serially diluted, spotted onto synthetic Histidine-dropout medium plates (SD-His) and SD-His plates with 0.1 *μ*g/mL FK506, and incubated at 30 °C for two days.

### The putative catalytic site of Fks1

Previous studies showed the GT-A fold adopts a conserved α/β/α sandwich with active site in the center (*33, 34*). Like other GT-A fold proteins, the cytosolic GT domain of Fks1 consists of a central nine-stranded β-sheet surrounded by two groups of α-helices. The putative active site of Fks1 was proposed to be around the end of central β-sheet, which is next to the transmembrane region. The putative active site features many charged residues, including E851, D1080, K1082, D1102, E1155, K1172, E1173, E1221, D1222, K1246, and R1248 (**Fig. 2b**). Among them, D1102 and D1222 correspond to the second and third aspartic acids of the conserved D, D, D signature in cellulose synthases (D180, D246, and D343 in BcsA). In the structure of BcsA, D246 in a DxD motif coordinates the divalent cation, whereas D343 in a TED motif of the finger helix is responsible for substrate binding and glucose polymer elongation(*26, 27, 31*). To reveal function of these key residues in detail, we superimposed the GT domain of Fks1 with the GT domain of bacterial BcsA and *Bacillus subtilis* SpsA (**Supplementary Fig. 7a-b**)(*26, 35*). We found their root mean square deviation (r.m.s.d.) are 2.4Å and 2.7Å respectively, indicating the GT domain of Fks1 is highly conserved. SpsA is a glycosyltransferase that transfers the glycosyl moiety from an activated UDP-sugar donor to a specific acceptor and was implicated in the synthesis of the spore coat of *Bacillus subtilis*. By investigating the substrates positions in the BcsA and SpsA structures and the active site of Fks1, we propose that E851, D1080, K1082, D1102, and N1104 at the top of the active site of Fks1 stabilize the sugar donor UDP-Glc, H1195, E1221, D1222, and K1246 probably form the acceptor glucose binding site, catalyze the sugar transfer, and participate in glucan elongation, and K1172 and E1173 in IF1, E1155, and R1248 may stabilize the glucan product in Fks1 (**Supplementary Fig. 7a-c**). All these residues in the active site are conserved among fungal 1,3-β-glucan synthases (**Supplementary Fig. 1**).

To evaluate the structure-function relationship of the key residues identified above, we performed in vivo growth complementation assays of the *fks1* mutants using the yeast *fks1Δ* strain. Previous studies indicated that the *fks1Δ* strain is hypersensitivity to the immunosuppressant FK506(*11*), and this deficiency can be partially complemented by a plasmid carrying the wild-type *fks1* gene. We found that E851L, D1080L, K1082L, D1221L, D1102L/N1104L, E1155L, K1172L/E1173L, H1195L, N1220L/E1221L, and K1246/R1248L mutants disrupted Fks1 function significantly as they were unable to rescue *fks1Δ* yeast growth (**Fig. 2c**).

The central catalytic region of our Fks1 structure in the apo-state is superimposable with the structure of BcsA for both the GT domain and transmembrane glucan-transporting domain (**Figs. 2a, 3a–d**). However, the GT domain of Fks1 is shifted upwards and away by ∼3–8 Å from the membrane region relative to that of BcsA (**Fig. 3b**). This structural feature leads to a more open active site, which probably facilitates substrate binding. We hypothesize that the GT domain will move downward upon substrate binding and thus push the glucan product out. To further confirm this hypothesis, we analyzed the cryo-EM structure of Fks1 using 3D variability analysis (3DVA), which is a newly developed software in CryoSPARC to reveal the continuous variability and discrete heterogeneity of a structure (*36*). We found that the cytosolic domain indeed moves up and down relative to the transmembrane region in two extremes, and accordingly the substrate binding site becomes more open and closed (**Supplementary Video 1**). We also noticed that a motif in GT domain (R862-V870) interacts a cytosolic loop between TMH15 and HH1 in CTD when the cytosolic domain moves down (**Supplementary Video 1, Supplementary Fig. 5a**). The CTD may regulate the activity of Fks1 through this interaction.

Interestingly, we found the predicted model by AlphaFold2 (*37, 38*) is more compact than our cryo-EM structure, although these two structures are superimposable with an r.m.s.d. of ∼2.0 Å. Compared to cryo-EM structure, the cytosolic domain and CTD in predicted model undergo rigid-body movement toward each other, forming a more closed active site (**Supplementary Fig. 8**). The predicted model of Fks1 by AlphaFold2 probably represents a substrate bound conformation.

### The putative transmembrane glucan transporting channel

The putative transmembrane glucan-transporting channel of Fks1 is proposed to be positioned directly below the active site and composed of TM5–10 (**Supplementary Fig. 9a, Fig. 3a-d**). Unlike the channel accommodating the translocating glucan in the BcsA and CesA structures(*26, 28*), there is no glucan observed in our structure of Fks1, indicating the structure to be in apo state. Furthermore, the putative glucan-transporting channel is only accessible from cytosolic side, but is closed in the extraplasmic side. Structural alignment between TM5–10 of Fks1 and corresponding region of BcsA and CesA8 revealed that TM5-7 and TM9-10 are well aligned with an r.m.s.d. of ∼3.0Å, while TM8 of Fks1 is shifted inward and occupy the glucan transporting path, resulting in a closed channel (**Fig. 3d, Supplementary Fig. 6**). Furthermore, compared with the ordered cytosolic entry of the channel in BcsA and CesA, which is mainly formed by three horizontal interface helices (IF1–3), a transmembrane helix (TM5 in BcsA and TM2 in CesA), a conserved finger helix and gating loop, the corresponding sequences of IF2–3 and the gating loop in Fks1 are flexible, and IF1 moves toward the channel by ∼5 Å (**Fig. 3c, d**). These structural features of Fks1 are likely caused by the absence of the glucan substrate. The AlphaFold2 predicted model of Fks1 showed IF2 and IF3 fold right under active site like in CesA structure, and may represent the substrate bound conformation of Fks1 (**Supplementary Fig. 8**). Although the flexible regions are not conserved between Fks1 and cellulose synthase, we found they are still essential for Fks1’s function according to the significant disruption of the K1261L mutation in IF2 and the IF3/gating loop deletion to Fks1 function in the growth complementation assay (**Fig. 2c**). Taken together, we propose that TM8 and IF1 need to move outward upon glucan synthesis to open the channel for glucan translocation, and IF2–3 play important roles during this process.

Crossing the membrane, the putative glucan-transporting channel featured with many hydrophobic residues and few polar residues (such as D684, N1300, N1301, S1307, T1314, T1359 and S1361) (**Supplementary Fig. 9a, b**). To investigated the functional significance of the polar residues, we introduced three mutations (N1300F/N1301F, T1314F, and S1361F) and performed complementation assays on *fks1*Δ cells. We found that all three mutations significantly disrupted Fks1 function *in vivo*, as they were unable to rescue the *fks1Δ* yeast growth than the plasmid carrying the WT *FKS1* (**Supplementary Fig. 10c**). Because the glucan is negatively charged, we suggest that the polar residues in the channel may assist the glucan transporting.

### Structural interpretation of echinocandin resistance

Previously reported echinocandin resistance mutations are mainly distributed in three “hot spot” regions (HS1: 635–649, HS2: 1354–1361, and HS3: 690–700). We found that these regions belong to TM5, TM8, and TM6 of Fks1, respectively, all of which are key TMs of the putative glucan transporting channel and distal from the active site (**Fig. 3e**). More specifically, HS1 and HS2 interact with each other, and this interface would be the major region to undergo conformational changes if TM8 shifts outward during glucan transport, as proposed above. We found two frequent echinocandins mutants, S643P and S643Y, in HS2 that maintained native Fks1 function in the growth complementation assay (**Fig. 3f**). Therefore, echinocandins likely inhibit Fks1 activity by affecting glucan translocation rather than synthesis.

### Two extracellular disulfide bonds of Fks1 are important for glycosyltransferase activity

The extracellular side of Fks1 only contains a few loops, but two disulfide bonds (Cys-658 with Cys-669 and Cys-1328 with Cys-1345) were found in the exoplasmic loops between TM5 and TM6 (EL3) and TM7 and TM8 (EL4) of Fks1, respectively (**Fig. 4a**). The two disulfide bonds are very close to neighboring transmembrane helices and appear to shorten EL3 and EL4. Given that TM8 may shift dramatically to open the channel during glucan transport as proposed above, the limited relative flexibility of TM5–8 by disulfide bonds may be important for regulating Fks1 activity.

**Fig. 4.**
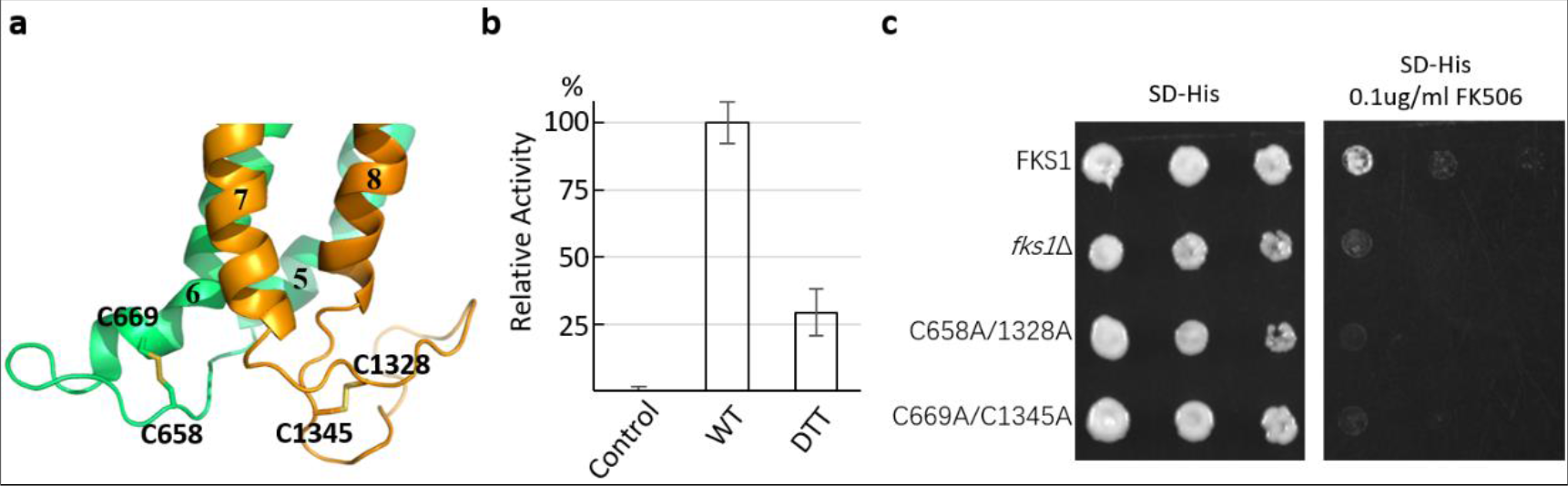
The two extracellular disulfide bonds of Fks1. **a** Two disulfide bonds (Cys-658 with Cys-669 and Cys-1328 with Cys-1345) are located in the exoplasmic loops between TM5 and TM6 (EL3), and TM7 and TM8 (EL4) of Fks1, respectively. **b** UDP-Glc hydrolysis activity of wild type (WT) Fks1 with or without DTT. All reactions were performed with Rho1 and GTPγS. The reaction without Fks1 was used as the control. Data points represent the mean ± SD in triplicate. **c** Growth complementation of *fks1*Δ cells with a plasmid carrying Fks1 mutant (C658A/C1328A or C669A/C1345A). The empty plasmid (*fks1*Δ) and a plasmid carrying wild-type Fks1 (FKS1) were used as controls.

We first performed the *in vitro* UDP-Glo glycosyltransferase assay to investigate the function of the disulfide bonds and found that Fks1 activity is reduced significantly by DTT, a disulfide bond reducing agent (**Fig. 4b**). Furthermore, we generated two FKS1 double mutants (C658A/C1328A and C669A/C1345A) and performed *in vivo* growth complementation assays of the mutants using the yeast *fks1Δ* strain. We found that neither C658A/C1328A nor C669A/C1345A mutants were able to rescue *fks1Δ* yeast growth on plates containing 0.1 *μ*g/mL FK506, supporting the essential functional role of the two disulfide bonds (**Fig. 4c**).

In addition, an N-glycan on Asn-1849 was found to locate directly above the disulfide bond formed between Cys-1328 and Cys-1345 and likely contributes to the stability of the structure (**Fig. 1c, Supplementary Fig. 4**).

## Discussion

As key enzymes in cell walls construction, Fks1 and cellulose synthases synthesize 1,3-β-glucan and 1,4-β-glucan (cellulose) respectively. Despite the low sequence identity of 11%–13%, the core domains of Fks1 and cellulose synthases from bacteria and plants share a conserved fold containing a cytosolic GT domain and a transmembrane glucan-transporting domain. This structural and functional similarities indicate that Fks1 functions by using a similar mechanism as cellulose synthases.

Remarkably, our structure of Fks1 was determined in the apo-state, while previous high-resolution structures of cellulose synthases were all reported in complex with substrate. Therefore, we proposed a working model of Fks1 based on the analysis of our structure and structural comparison with cellulose synthases (**Fig. 5**). In the apo-state of Fks1, the putative active site is open for substrate binding, whereas the putative glucan transporting channel is closed and occupied by TM8. Flexible IF2–3 probably also facilitate substrate entering the active site. We further propose that substrate binding and the glycosyl transfer reaction trigger the movement of the GT domain toward the glucan transporting channel and thus lead to channel opening and glucan elongation. This working model is also supported by the Fks1 structure predicted by AlphaFold2 and the 3DVA analysis of our cryo-EM structure of Fks1 as mentioned above. Capturing Fks1 in complex with substrates will undoubtedly provide more insights into this model.

**Fig. 5.**
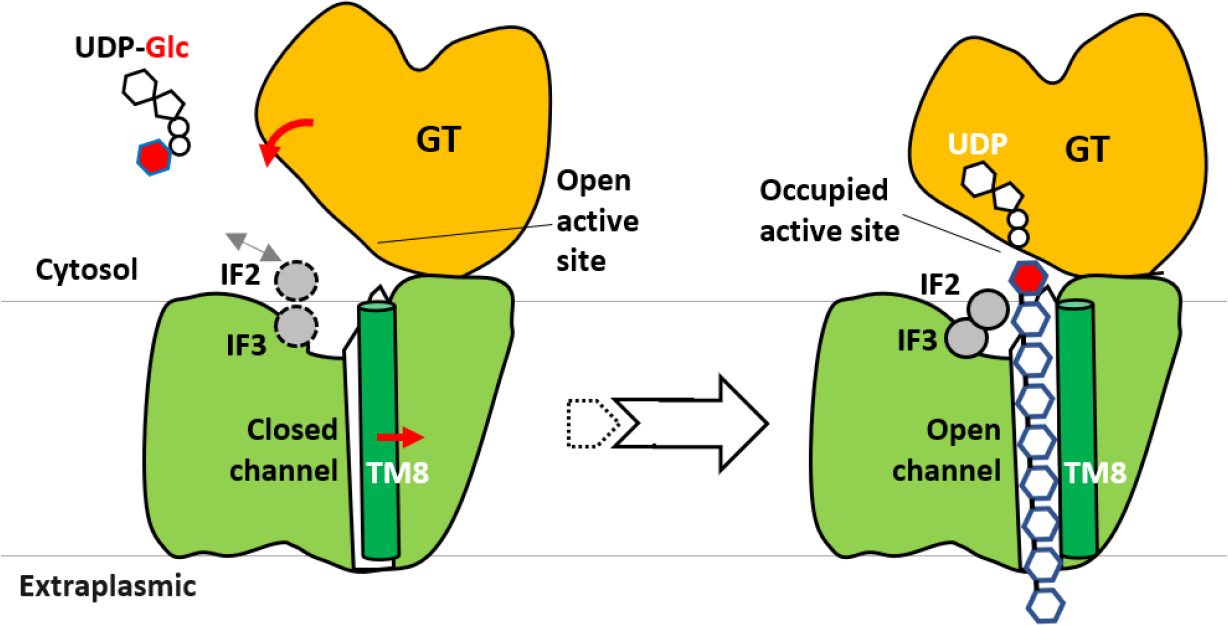
Proposed catalytic mechanism of Fks1. The cryo-EM structure reflects the apo-state (left) with flexible IF2–3 and an empty open, active site for substrate binding. The glucan transporting channel is closed and engaged by TM8. Substrate binding and subsequent reaction may trigger the downward movement of the GT domain, which pushes the glucan product out of the cell. IF2–3 may stabilizes the glucan during glucan elongation, and TM8 is shifted outward, resulting in an open channel. The conformation of Fks1 in complex with substrates (right) is modeled according to the structure of cellulose synthases in complex with UDP and cellulose. More details are provided in the text.

A recent study reported a preliminary hexameric structure of the putative *Candida glabrata* 1,3-β-glucan synthase (GS) complex at ∼14 Å resolution using cryo-electron tomography (cryo-ET) and subtomogram analysis (**Supplementary Fig. 10a**) (*39*). To investigate the mechanism of Fks1 in vivo, we compared our high resolution Fks1 structure with the cryo-ET map of the hexameric GS (**Supplementary Fig. 10a, b**). Surprisingly, we found our Fks1 structure is different from one subunit of the cryo-ET map in terms of either size or features. Specifically, the transmembrane region of our structure is much larger than one subunit of the cryo-ET map, while the cytosolic region of our structure is much shorter. In the cryo-ET study, the authors proposed that each subunit of the putative GS complexes contains two notable cytosolic domains, the N-terminal and central catalytic domains. However, our structure of Fks1 only contains one large cytosolic GT domain.

Moreover, we found the cryo-ET map of putative hexameric GS is well superposed with the structure of the hexameric Pma1, which is the most abundant protein in the plasma membrane of most fungi including *Candida glabrata* (**Supplementary Fig. 10c**) (*40*). Notably, the cytosolic domains of Pma1 are well consistent with features of the cryo-ET map. Assignment of particles in low resolution cryo-electron tomography is always challenging. Although the authors compared native and Fks1 overexpressing strains to prove the identity of particles to be GS in that cryo-ET study, the evidence was not direct and was easily affected by bias. The possibility that the most abundant hexameric Pma1 in plasma membrane was mistaken for the GS can’t be ruled out. Therefore, whether the reported cryo-ET structure of putative hexamric GS is corresponding to GS or Pma1 still needs more studies to confirm. According to our results, we tend to believe that Fks1 functions as a monomer, rather than a hexamer.

In addition, the inhibitory mechanism of echinocandins and ibrexafungerp and the mechanism of echinocandin resistance mutations remain unclear. Capturing structures of Fks1 in complex with inhibitors will help to unveil these mechanisms. Further research is required to fully understand the regulatory role of Rho1 toward Fks1 and the functional roles of the N- and C-terminal domains of Fks1. Besides, the invisible N-terminal loop of Fks1 is also probably essential as we found that the deletion of first 140 residues strongly decreased Fks1’s function in the growth complementation assay (**Fig. 2c**). How this loop works also awaits further studies.

In conclusion, our cryo-EM structure and structure-based mutagenesis analysis of Fks1 have provided deep insights into the molecular mechanisms for glycosyltransferase activity and transmembrane glucan transport of a fungal 1,3-β-glucan synthase and echinocandins resistance by Fks1 mutants. Given the critical role of 1,3-β-glucan synthase in pathogenic fungal infections, our work provides a platform for developing new small anti-fungal molecules.

## Materials and Methods

### Expression and purification of Fks1

We tagged *FKS1* with a C-terminal triple-FLAG to the modified pRS423 vector. The yeast strain BY4742 was transformed and cultured by synthetic Histidine-dropout medium (SD-His) for about 20h and then transferred to YPG medium (10g Yeast extract, 20g Peptone, 20g D-Galactose per liter) for 12 hours before harvest. Cells were resuspended in lysis buffer (20 mM Tris-HCl, pH 7.4, 0.2 M sorbitol, 50 mM potassium acetate, 2 mM EDTA, and 1 mM phenylmethylsulfonylfluoride (PMSF) and then lysed using a French press at 15,000 psi. We centrifuged the lysate at 10,000 ×g for 30 min at 4 °C, and collected the supernatant for another centrifuge cycle at 100,000 ×g for 60 min at 4 °C. The membrane pellet was collected and then resuspended in buffer A containing 10% glycerol, 20 mM Tris-HCl (pH 7.4), 1% DDM, 0.1% cholesteryl hydrogen succinate (CHS), 500 mM NaCl, 1 mM MgCl2, 1 mM EDTA, and 1 mM PMSF. After incubation for 30 min at 4 °C, the mixture was centrifuged for 30 min at 100,000 ×g to remove the insoluble membrane. We loaded the supernatant into a pre-equilibrated anti-FLAG (M2) affinity column (GenScript) at 4 °C and washed the affinity gel with buffer B (20 mM HEPES, pH 7.4, 150 mM NaCl, 0.01% lauryl maltose neopentyl glycol (LMNG), 0.001% CHS, and 1 mM MgCl_2_). The proteins were eluted with buffer B containing 0.15 mg/mL 3 × FLAG peptide and were further purified in a Superose 6 10/300 Increase gel filtration column in buffer C (20 mM HEPES, pH 7.4, 150 mM NaCl, 0.003% LMNG, 0.0003% CHS, and 1 mM MgCl2). The purified proteins were assessed by SDS-PAGE gel and concentrated for cryo-EM analysis. To maintain consistency with previous functional assays(*17, 18, 25*), the samples used for UDP-Glo glycosyltransferase assay were using the same procedure as described above except that CHAPS and CHS were used to solubilize the membrane and stabilize the membrane protein.

### Expression and purification of Rho1

*RHO1* was cloned into pET-28a vector with a C-terminal His tag and expressed in E. coli BL21(DE3). Bacteria were grown in Luria-Bertani (LB) broth at 37°C until the optical density at 600 nm (OD600) reached approximately 0.6, and then cells were induced with 1mM isopropyl-β-d-1-thiogalactopyranoside (IPTG) at 25°C overnight. The cell pellet was collected and re-suspended in 50mL of buffer (20 mM HEPES (pH 7.4), 150 mM NaCl, 1 mM MgCl2, 1 mM MnCl2 and 1 mM EDTA) supplemented with protease inhibitors (1 mg/ml each of DNase, pepstatin, leupeptin, and aprotinin and 1 mM PMSF). Cells were lysed on ice by sonication. Pellets were removed by centrifugation for 30 min at 12,000 rpm, and the supernatant was loaded onto a 2ml column of Nickel-NTA agarose. The column was washed with 20 mM HEPES pH 7.4, 20 mM imidazole, 150 mM NaCl, and Rho1 was eluted with wash buffer supplemented with 200 mM imidazole. The protein eluate was concentrated and further purified by size-exclusion chromatography on a Superose6 10/300 GL column (GE Healthcare). Peak fractions were collected for functional assay.

### Cryo-electron microscopy

2.5μL aliquots of Fks1 at a concentration of about 3 mg/mL were placed on glow-discharged holey carbon grids (Quantifoil Au R1.2/1.3, 300 mesh) and were flash-frozen in liquid ethane using a FEI Vitrobot Mark IV. Grids were screened in in a 300-keV FEI TF30 electron microscope from School of Basic Medical Sciences, Peking University. Cryo-EM data was collected automatically with SerialEM in a 300-keV FEI Titan Krios electron microscope from School of Physics, Peking University and Institute of Biophysics Chinese Academy of Sciences. We collected Cryo-EM data with defocus values ranging from -1.0 to 2.0μm. The microscope was operated with a K3 direct detector at a nominal magnification of 130,000×. The total doses were 50-60 electrons per Å^2^ at the sample level.

### Cryo-EM image processing

We used the program MotionCorr-2.0 (*41*)^5^ for motion correction, and CTFFIND-4.1 (*42*)^5^ for calculation of contrast transfer function parameters. We used RELION-3 for particle picking and extraction, and used CryoSPARC for all remaining steps (one cycle of 2D classification, one cycle of ab-inito reconstruction, one cycle of heterogeneous refinement, one cycle of CTF refinement, and one cycle of local refinement)) (*36, 43*). The resolution of the map was estimated by the gold-standard Fourier shell correlation at a correlation cutoff value of 0.143.

We totally collected 1670 raw movie micrographs. A total of 1,077,552 particles were picked automatically for 2D and 3D classifications. Based on the quality of the three 3D classes, 184,415 particles were retained for further refinement, resulting in a 3.0-Å average resolution 3D map using C1 symmetry.

### Structural modeling, refinement, and validation

We used the predicted Fks1 structure by Alphafold2 as initial model (*37, 38*), fitted it into our map, and corrected it in COOT (*44*) and Chimera (*45*). The complete Fks1 model was refined by real-space refinement in the PHENIX program (*46*) and subsequently adjusted manually in COOT. Finally, the model was validated using MolProbity (*47*). Structural figures were prepared in Chimera and PyMOL (https://pymol.org/2/).

### UDP-Glo Glycosyltransferase Assay

The activity of WT Fks1 were measured using the UDP-Glo™ Glycosyltransferase Assay from Promega (Catalog: V6961), which can measure the activity of any glycosyltransferase (GT) that uses a UDP-sugar as a substrate. Each reaction contained 0.2 μg purified Fks1, 20 mM HEPES, pH 7.4, 150 mM NaCl, 0.6% CHAPS, 0.06% CHS, 10% glycerol, and 1mM EDTA in a total volume of 5 μL. For other reaction conditions, 0.05μg purified Rho1, 1mM GTP-γS, 2mM glucose, 2mM Mg^2+^, 1mM Caspofungin, or 1mM Enfumafungin was also added. The mixture was first incubated for 30 min at 37 °C, and then 5 μL of UDP Detection Reagent was added. The mixture was incubated for 60 min at room temperature, and then luminescence was detected by Synergy H1 Hybrid Multi-Mode Microplate Reader (BioTek).

### Colony growth assay

Wide type yeast strain and the *FKS1* knockout strains were first grown in SD medium at 30°C overnight to the same OD. Then 1:10 serial dilutions of the cells were spotted onto SD plates with or without 0.1 μg/mL FK506, incubated at 30°C for 2 d, and examined for growth.

## Acknowledgements

Cryo-EM data were collected in the Electron Microscopy Laboratory of Peking University, and Cryo-EM Platform at the Center for Biological Imaging (CBI, cbi.ibp.ac.cn) at the Institute of Biophysics, Chinese Academy of Sciences. We thank Xuemei Li, Zhenxi Guo, Boling Zhu, Xujing Li, and Xiaojun Huang for facilitating data collection. We thank Huichao Deng and Xiangming Wang in Institute of Biophysics for helps in construct preparation. We thank Liwen Bianji (Edanz) for editing the English text of a draft of this manuscript. This work was supported by grants from the National Natural Science Foundation of China (32171212 to L.B., No. 32071207 to C.Y., 82071658 to J.H.), Peking University (to L.B.), the National Basic Research Program of China (973 Program, No. 2012CB917202 to C.Y.), the Beijing Nova Program (Z201100006820010 to J.H.), and National Key R&D Program of China (2018YFC1004403 to J.H.).

## Author Contributions

C.Z., D.C., Z.Y., J.H., J.Q., C.Y., and L.B. conceived and designed the experiments. C.Z., D.C., Z.Y., Z.W., J.H., P.Z., and L.B. performed the experiments. C.Z., D.C., Z.Y., J.H., J.Q., C.Y., and L.B. analyzed the data. J.H., J.Q., C.Y., and L.B. wrote the manuscript with input from all authors.

## Data Availability

The cryo-EM 3D map and the corresponding atomic model of the Fks1 have been deposited at the EMDB database and the RCSB PDB with the respective accession codes of EMD-XXXX and YYYY.

---

**Supplemental Table 1.** Cryo-EM data collection, refinement, and validations.

|  |  |
| --- | --- |
| Sample | *S. cerevisiae* Fks1 |
| Map and Model | EMD-XXXX, PDB ID: YYYY |
| **Data collection and processing** |  |
| Microscope | FEI Titan Krios |
| Voltage (kV) | 300 |
| Electron exposure (e<sup>-</sup>/Å<sup>2</sup>) | 50 |
| Defocus range (-μm) | 1.0-2.0 |
| Pixel size (Å) | 1.0552 |
| Micrographs (no.) | 1670 |
| Initial particle images (no.) | 1,077,552 |
| Final particle images (no.) | 184,415 |
| Symmetry imposed | C1 |
| Map resolution (Å) | 3.0 |
| FSC threshold | 0.143 |
| Map resolution range (Å) | 3.0 - 250 |
| **Refinement** |  |
| Map sharpening *B* factor (Å<sup>2</sup>) | 124.1 |
| Model composition |  |
| Non-hydrogen atoms | 12,183 |
| Protein residues | 1489 |
| R.m.s. deviations |  |
| Bond lengths (Å) | 0.01 |
| Bond angles (°) | 0.90 |
| Validation |  |
| MolProbity score | 1.99 |
| Clash score | 11.2 |
| Poor rotamers (%) | 0 |
| Ramachandran plot |  |
| Favored (%) | 93.29 |
| Allowed (%) | 6.71 |
| Disallowed (%) | 0 |

