## Supplemental Figures for "Structure of a fungal 1,3-β-glucan synthase"

### Supplemental Figures and Video

**Supplementary Video 1.** 3DVA analysis of the Fks1 structure, revealing the movement of the soluble glycosyltransferase domain (GTD) and C-terminal domain (CTD) relative to the transmembrane region of Fks1.

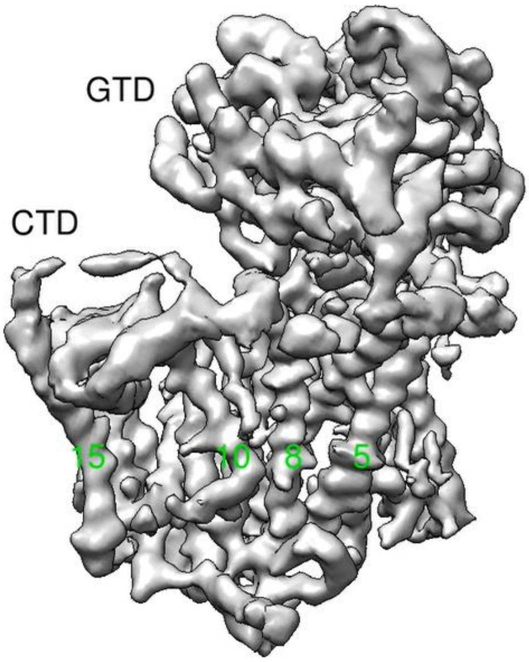

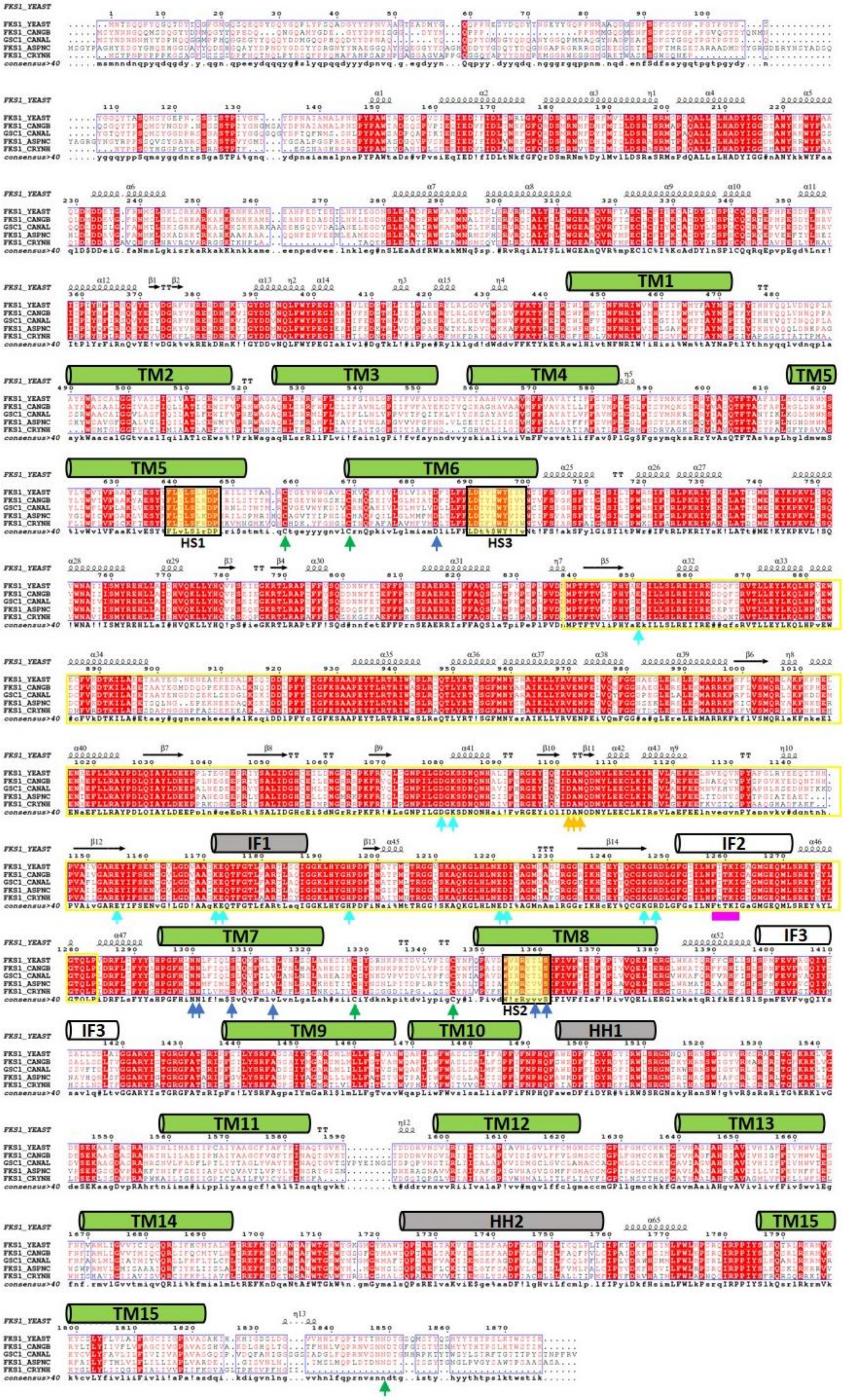
FKS1_YEAST
FKS1_CANGB
GSC1_CANAL
FKS1_ASPNC
FKS1_CRYNH
consensus>40
TM1
TM2
TM3
TM4
TM5
TM6
TM7
TM8
TM9
TM10
TM11
TM12
TM13
TM14
TM15
HS1
HS2
HS3
IF1
IF2
IF3
HH1
HH2

**Supplemental Fig. 1 Sequence alignment of Fks1 from different species.**

YEAST: *Saccharomyces cerevisiae*, CANGB: *Candida glabrata*, CANAL: *Candida albicans*, ASPNC: *Aspergillus niger*, CRYNH: *Cryptococcus neoformans*. Transmembrane helices are highlighted by green cylinders according to our Fks1 structure. The GT domain sequence inserted between TM6 and TM7 is highlighted by a yellow rectangle. Red arrows indicate key residues in the active site. An orange arrow indicates the metal-coordinating DAN motif. A yellow background highlights hot spots 1–3 (HS1–3). Green arrows highlight two disulfide bonds. Interface helices (IF1–3) and horizontal helices (HH1–2) are highlighted by black or grey cylinders.

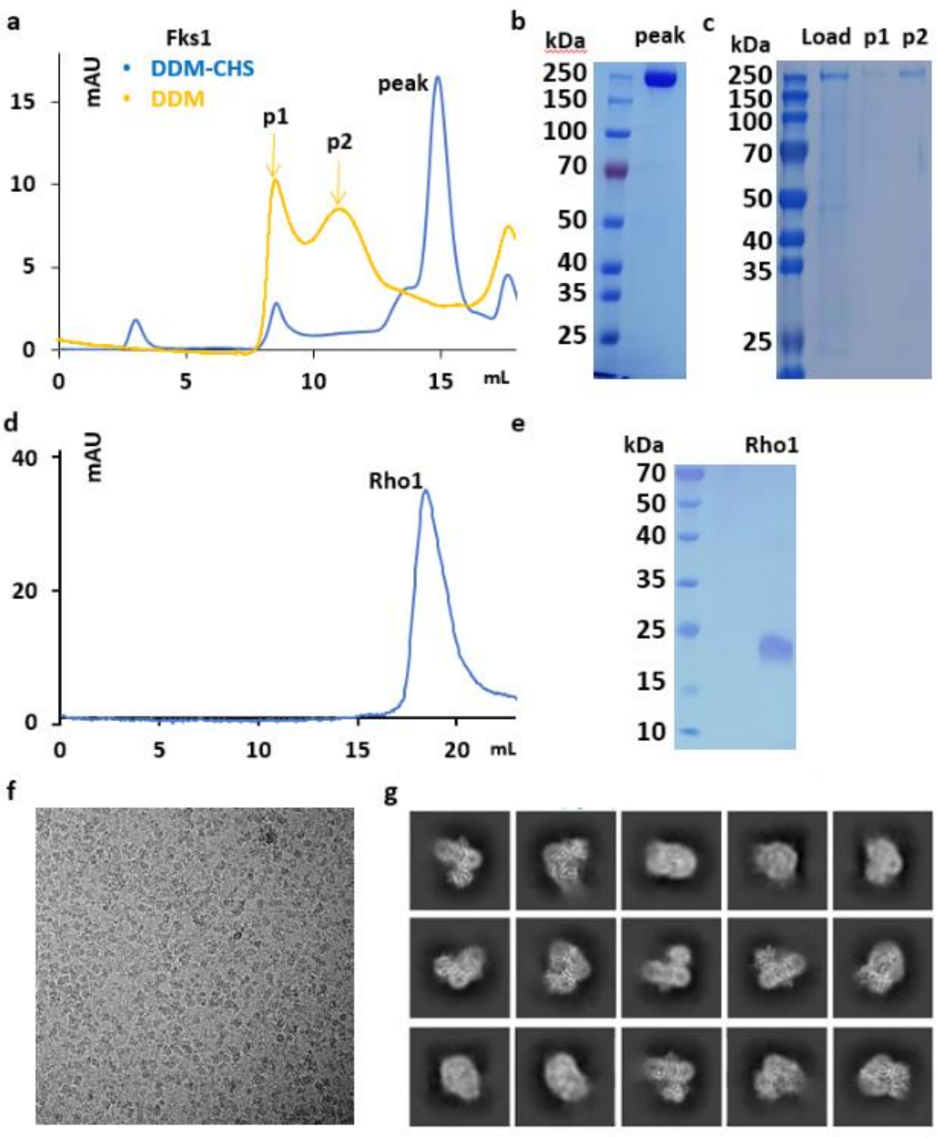


**Supplemental Fig. 2 Purification and analysis of Fks1 and Rho1. a** Gel filtration profile of Fks1 purified with CHS (blue) or without CHS (orange). **b** Coomassie blue-stained SDS-PAGE gel of the peak fraction of Fks1 purified with CHS. **c** Coomassie blue-stained SDS-PAGE gel of the peak fractions of Fks1 purified without CHS. **d** Gel filtration profile of Rho1. **e** Coomassie blue-stained SDS-PAGE gel of the peak fraction of Rho1. **f** A representative electron micrograph. **g** Selected reference-free 2D class averages.

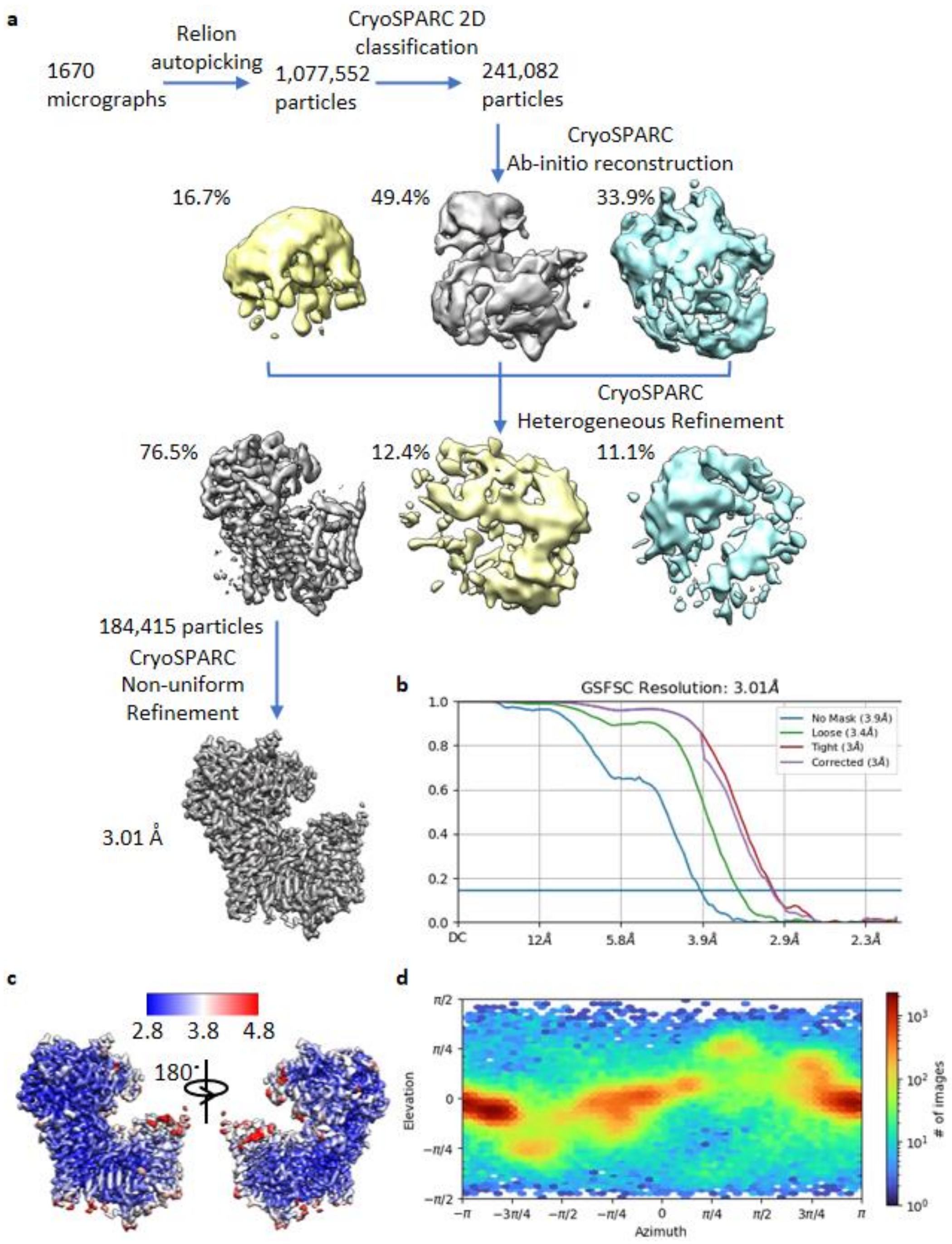


**Supplemental Fig. 3 Cryo-EM data processing and resolution estimation of Fks1. a** Cryo-EM data processing procedure. **b** Gold-standard Fourier shell correlation of two independent half 3D maps of Fks1. **c** Local resolution map of the 3.0-Å 3D map of Fks1. **d** Angular distribution of raw particles used in CryoSPARC 3D reconstruction of the 3.0-Å 3D map of Fks1.

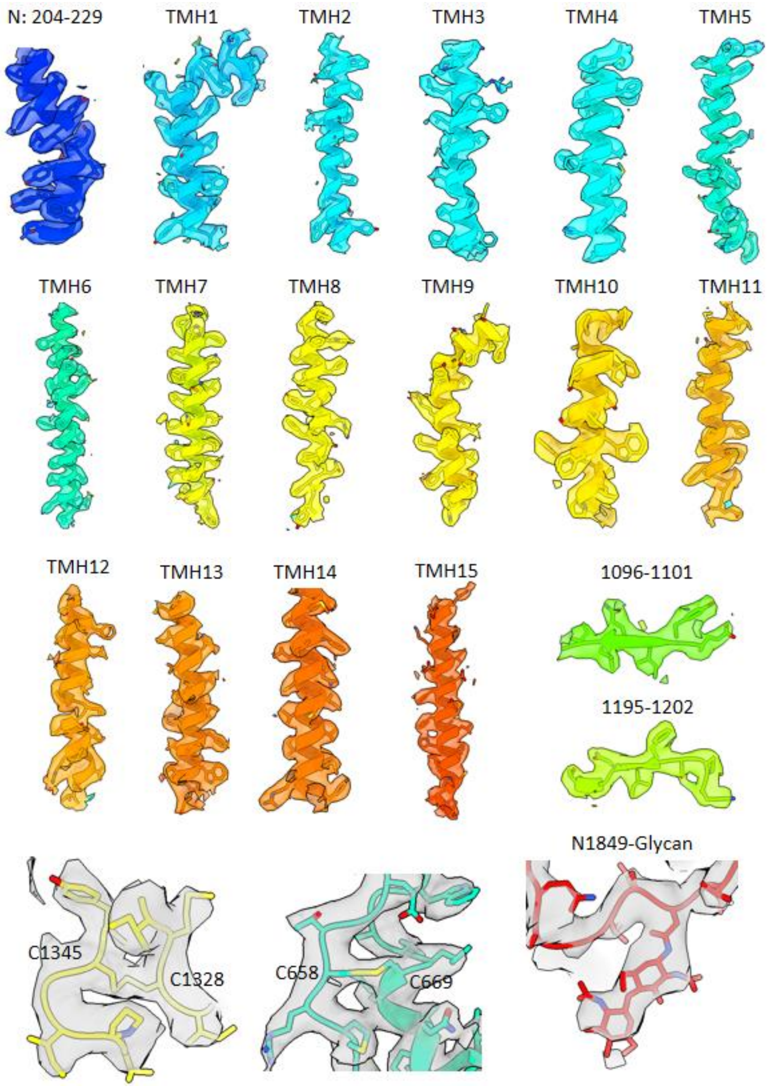


**Supplemental Fig. 4 Selected regions in the 3D map of Fks1 at 3.0 Å resolution superimposed onto the atomic model.**

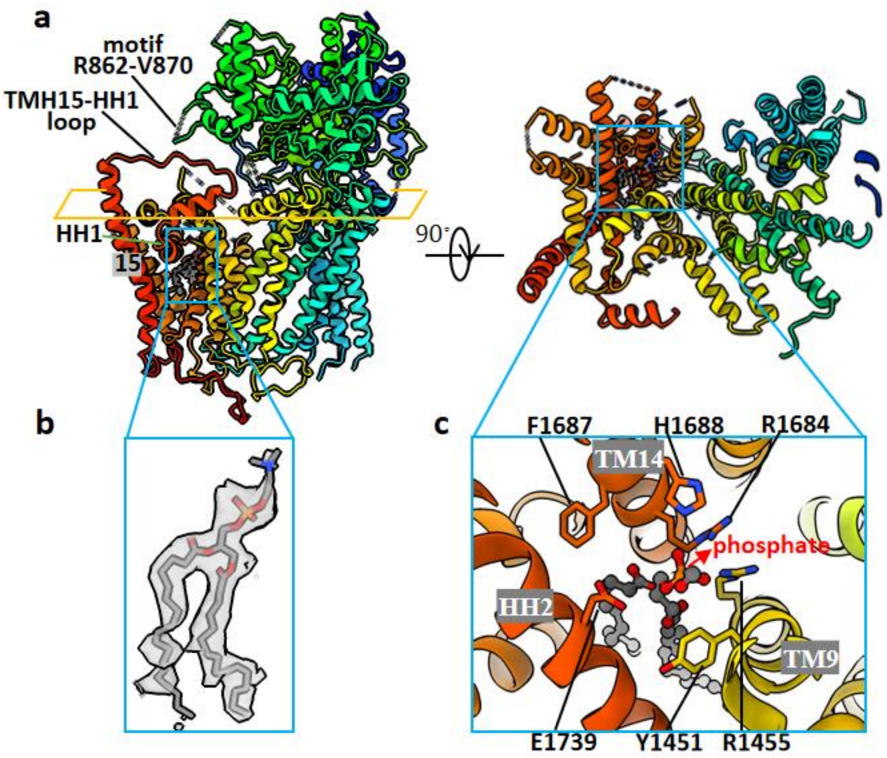


**Supplemental Fig. 5 An ordered lipid filling a cavity surrounded by TM9, TM14 and HH2. a** The overall structure of Fks1 is shown in a side view (left) and a top (cytosol) cut-in view (right). A motif in GT domain (R862-V870) is adjacent to the cytosolic loop between TMH15 and HH1 in CTD. **b** Electron density of the lipid. **c** A close-up view of the lipid interactions with TM9, TM14 and HH2.

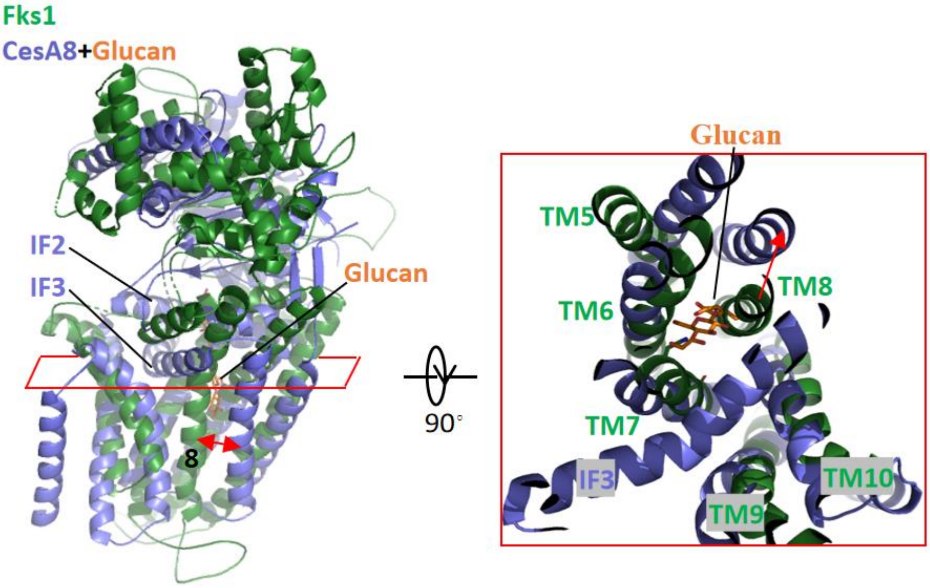


**Supplemental Fig. 6 Structural comparison between the central catalytic region of Fks1 (green) and CesA8 (PDB ID: 6WLB, blue) in complex with cellulose (orange) by aligning their respective TMs.** Compared with CesA8, TM8 of Fks1 is shifted inward and occupy the glucan transporting path, resulting in a closed channel.

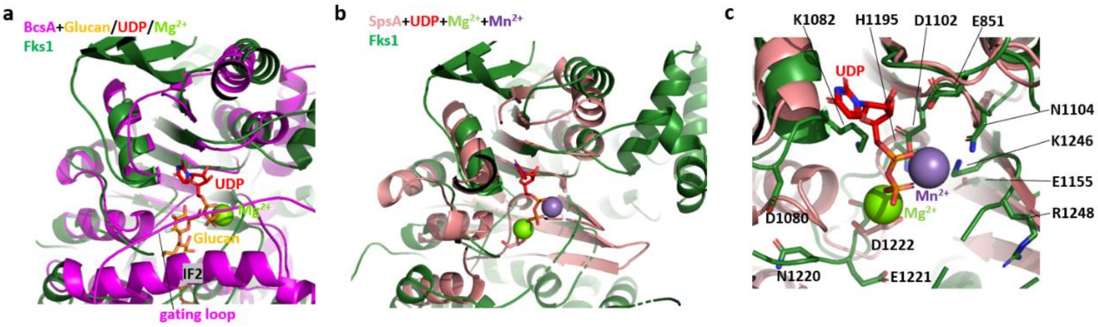


**Supplemental Fig. 7 Structural comparison between the soluble glycosyltransferase domains of Fks1, BcsA and SpsA. a** Structural comparison between the soluble glycosyltransferase domains of Fks1 (green) and BcsA (magenta) in complex with cellulose (orange), UDP (red), and $Mg^{2+}$ (lemon). **b** Structural comparison between the soluble glycosyltransferase domains of Fks1 (green) and SpsA (PDB ID: 1QGQ, salmon) in complex with UDP (red), $Mg^{2+}$ (lemon), and $Mn^{2+}$ (purple). **c** Zoom in view of the putative active site in panel **b**.

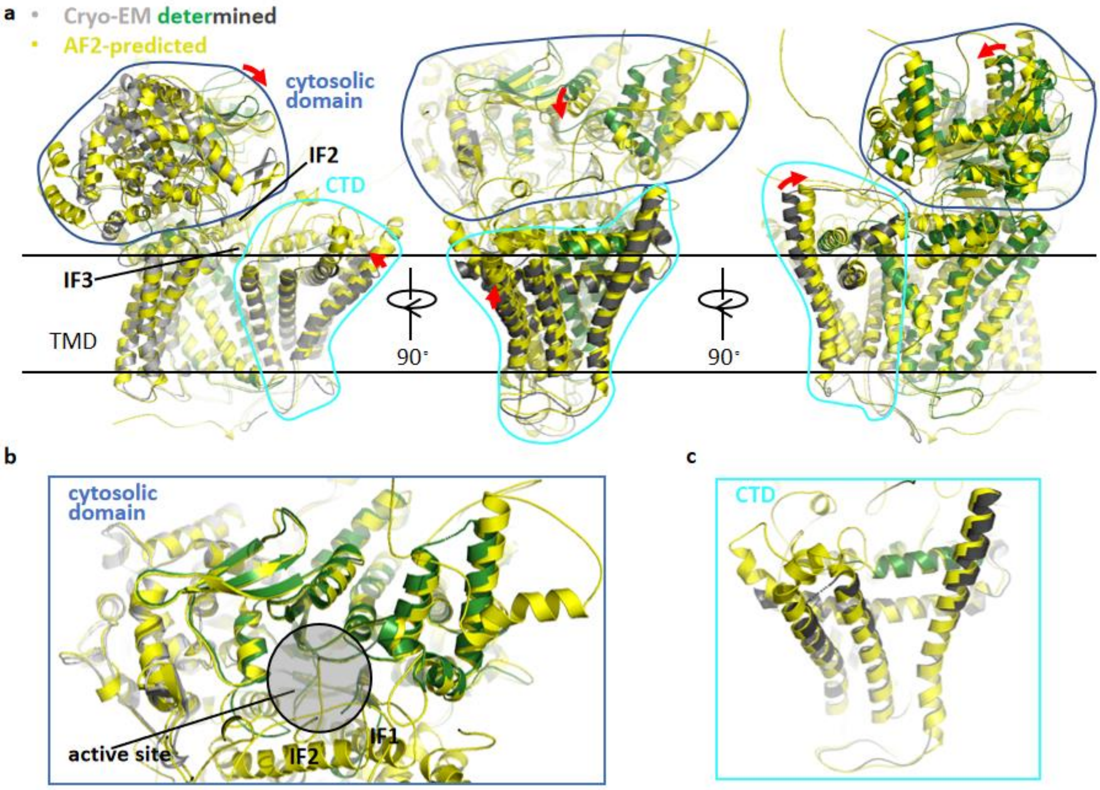


**Supplemental Fig. 8 Structural comparison between cryo-EM structure and AlphaFold2 predicted structure of Fks1. a** Two structures are superimposed by aligning TMH1-10 of Fks1. The predicted model is shown in yellow cartoon. The cryo-EM structure of Fks1 is split into three domains: the central catalytic region (green), N-terminal domain (grey), and C-terminal domain (CTD, black). The soluble cytosolic domain and CTD are outlined by blue and cyan lines respectively. The curved red arrows show the domain rotations between the two structures. **b** Superimposed cytosolic domain of two structures in the same view as middle figure in panel a. **b** Superimposed CTD of two structures in the same view as middle figure in panel a.

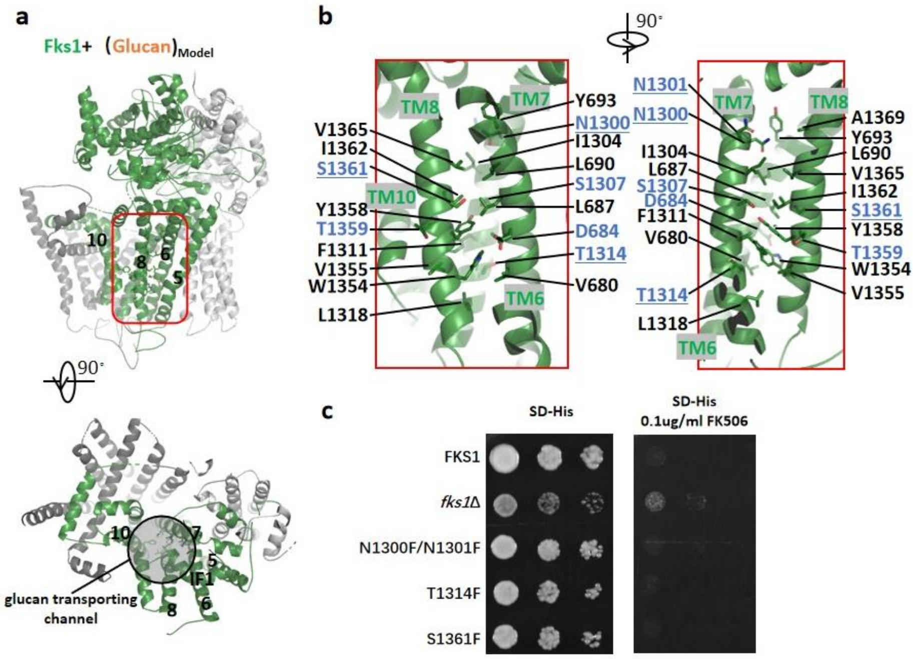


**Supplemental Fig. 9 Putative glucan transporting channel of Fks1. a** Side and top views of overall structure of Fks1 in cartoon representation. The central catalytic region (green) is sandwiched by N-terminal (grey) and C-terminal (black) regulatory domains. The putative glucan transporting channel is highlighted by a red rectangle in side view and a black circle in top view. **b** Two views of putative glucan transporting channel with residues forming the channel shown in stick representation. Labels of polar residue are highlighted by blue. Four underline labeled residues are selected for growth complementation assay. **c** Growth complementation of *fks1*Δ cells with empty plasmid (*fks1*Δ), or plasmid carrying either wild-type Fks1 (FKS1) or mutants. Cells were serially diluted, spotted onto synthetic Histidine-dropout medium plates (SD-His) and SD-His plates with 0.1 μg/mL FK506, and incubated at 30 °C for two days.

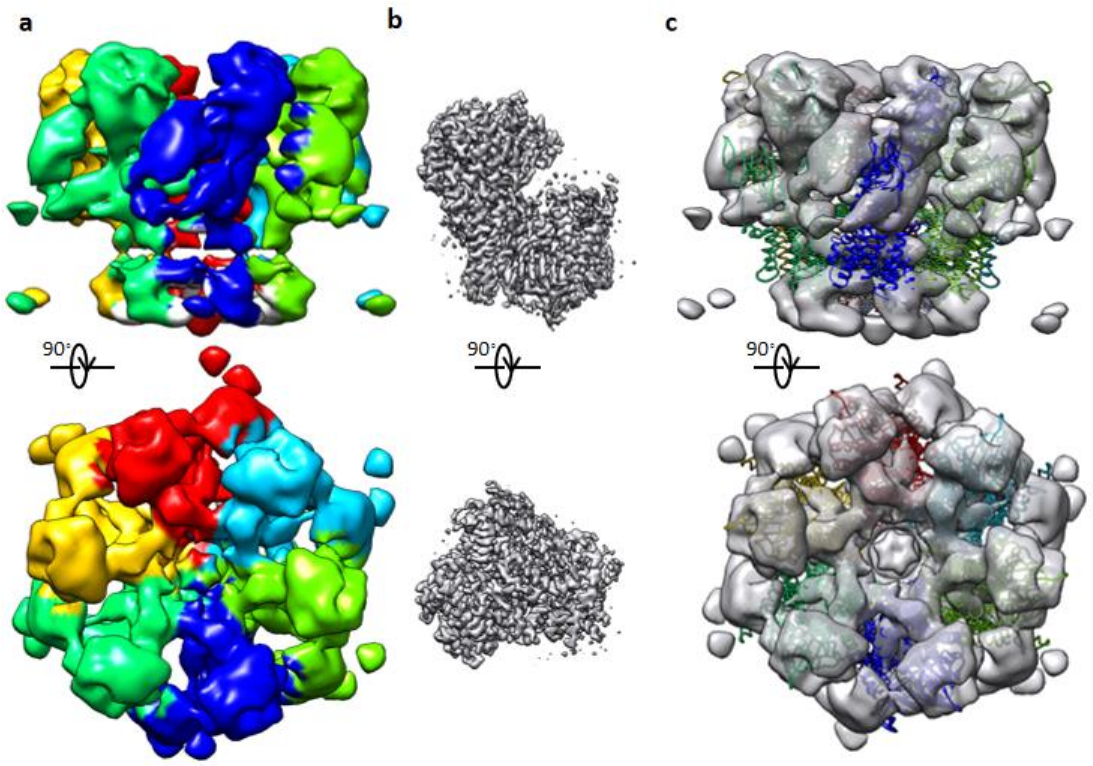


**Supplemental Fig. 10 Cryo-ET map of the putative *Candida glabrata* 1,3-β-glucan synthase. a** Overall cryo-ET map of the putative Candida glabrata 1,3-β-glucan synthase at ~14 Å resolution (EMD-23123). Six segmented subunits are shown in different colors. **b** Overall cryo-EM map of our Fks1 structure at ~3 Å resolution in the same view as figures in panel a. **c** The cryo-ET map superimposed with hexameric Pma1 structure (PDB: 7VH6) in the same view as figures in panel a.
